# Performance of GPT-4 on the American College of Radiology In-Service Examination

**DOI:** 10.1101/2024.02.15.580546

**Authors:** David L. Payne, Kush Purohit, Walter Morales Borrero, Katherine Chung, Max Hao, Mutshipay Mpoy, Michael Jin, Prateek Prasanna, Virginia Hill

**Affiliations:** Stony Brook University Hospital; Stony Brook University; Northwestern University Feinberg School of Medicine

## Abstract

**Objectives:** No study has evaluated the ability of ChatGPT-4 to answer image-rich diagnostic radiology board exam questions or assessed for model drift in GPT-4’s image interpretation abilities. In our study we evaluate GPT-4’s performance on the American College of Radiology (ACR) 2022 Diagnostic Radiology In-Training Examination (DXIT).

**Methods:** Questions were sequentially input into GPT-4 with a standardized prompt. Each answer was recorded and overall accuracy was calculated, as was logic-adjusted accuracy, and accuracy on image-based questions. This experiment was repeated several months later to assess for model drift.

**Results:** GPT-4 achieved 58.5% overall accuracy, lower than the PGY-3 average (61.9%) but higher than the PGY-2 average (52.8%). Adjusted accuracy was 52.8%. GPT-4 showed significantly higher (p = 0.012) confidence for correct answers (87.1%) compared to incorrect (84.0%). Performance on image-based questions was notably poorer (p < 0.001) at 45.4% compared to text-only questions (80.0%), with adjusted accuracy for image questions of 36.4%.

When the questions were repeated, GPT-4 chose a different answer 25.5% of the time and there was a small but insignificant decrease in accuracy.

**Discussion:** GPT-4 performed between PGY-2 and PGY-3 levels on the 2022 DXIT, but significantly poorer on image-based questions, and with large variability in answer choices across time points. This study underscores the potential and risks of using minimally-prompted general AI models in interpreting radiologic images as a diagnostic tool. Implementers of general AI radiology systems should exercise caution given the possibility of spurious yet confident responses.

## Introduction

Since its release to the public in November of 2022, ChatGPT, the flagship product of OpenAI (San Francisco, CA), has gained immense popularity, with reportedly 180.5 million monthly users as of February 2024 [1, 2]. The most current offering from OpenAI, GPT-4, is a large multi-modal artificial intelligence (AI) model capable of accepting both text and image prompts and outputting detailed responses in a wide variety of settings, and has demonstrated impressive performance on a variety of professional and academic tasks including high school AP examinations, the Bar exam, and notably for physicians, the USMLE exams [3]. GPT-4 has been able to achieve these results without the need for fine tuning, and has demonstrated superior performance compared to competitor models fine-tuned specifically for medical topics such as Med-PaLM [4].

Radiology is far at the forefront of the medical field in the development, implementation, and validation of AI tools [5]. While these technologies hold great promise for improving the accuracy and efficiency of diagnostic radiology (DR) while decreasing radiologist cognitive load, major concerns remain around the deployment of increasingly powerful AI image interpretation tools [6]. These include risks to patient safety, AI bias, data security, model drift, and disruption of the radiology workforce [7].

Many of the current radiology AI tools perform narrow tasks such as detection of intracranial hemorrhage, pulmonary embolism, breast cancer detection, spinal cord compression, detection of pulmonary nodules, as well as clinical decision support [8, 9]. However, given the rapid ascent of general AI models with multi-modality capabilities, it is important to assess the ability of these cutting edge and publicly available general models to perform on general radiology tasks including multi-modal image interpretation, image quality control, and screening guidelines.

Several studies have been carried out demonstrating impressive performance of ChatGPT on radiology board exam style questions, including questions drawn from or simulating the United Kingdom, Brazilian, Japanese, Canadian, and American board examinations [10, 11, 12, 13]. Additional work has shown the ability of ChatGPT to answer *Diagnosis Please* cases from *Radiology*, albeit with less impressive performance compared to percent correct on boards-style questions, unsurprising given the difficult nature of *Diagnosis Please* cases [14].

Within the above described studies however, the ability of ChatGPT to answer radiology questions was based solely on unimodal, i.e., text-only prompts, with diagnostic radiologists providing text description of radiology images to the model or excluding those questions altogether. *Li et al* also showed statistically significant model drift with worsening performance when testing ChatGPT several months apart [14]. Model drift within ChatGPT has also been observed across domains, and while the reason for these changes is being actively investigated, *Chen et al* have proposed that training on new data can have unexpected effects on other aspects of model performance [15].

To our knowledge, no previous work has evaluated the ability of GPT4 to answer image-rich DR board-style questions or tested ChatGPT for model drift in its ability for medical image interpretation. The purpose of this study is to assess the performance of GPT-4 on the DR In-Training Examination (DXIT), a yearly standardized examination prepared by the American College of Radiology (ACR) covering a wide breadth of DR topics and shown in several studies to be predictive of performance on the American Board of Radiology (ABR) Core Examination [16, 17, 18, 19].

## Methods

Checklist for Artificial Intelligence in Medical Imaging (CLAIM) was used as a guide in the execution of this study and in the preparation of this manuscript [20].

## Data Entry

A paid subscription was obtained for Chat GPT Plus, which includes access to GPT4, including vision capabilities.

The most current set of publicly available DXIT sample questions (from 2022) were retrieved from the ACR website, and permission to use these questions was obtained from ACR staff [21]. Data pertaining to resident performance on this question set, separated by topic and post graduate year (PGY) were also given by ACR staff.

A standardized prompt was given to GPT4, stating the following:

> *“Pretend you are a radiology resident. I am going to input a series of multiple choice radiology in-service examination questions for you to answer. Some questions are text-based and some include images. Answer these questions to the best of your ability. No harm will come to any patients based on your answers, as this is only a practice exam. In addition, give a brief explanation (less than 4 sentences) of your rationale, as well as a numerical value from 1-100 of your confidence in the correctness of your answers. Does this make sense?”*

Once an affirmative response was given by GPT4, screenshots of each question, including text, images, and multiple choice answer choices were taken, then pasted into the GPT4 user interface.

## Data Collection and Analysis

Question subject (neuroradiology, musculoskeletal, gastrointestinal, breast, etc.), question type (text-only vs. image-containing), correct response, GPT4 response, and level of reported confidence by GPT were recorded. Note was made for each question whether each selected answer by GPT was correct based on the official answer key included in DXIT. Additionally, the rationale for each answer choice was noted, and if there was a significant logical flaw in a correct answer (Figure 1), this was also recorded. These tasks, including determination of significant logical flaws, were performed by PGY-4 and PGY-5 radiology residents and performed from November 15th through December 11th 2023.

**Figure 1).**
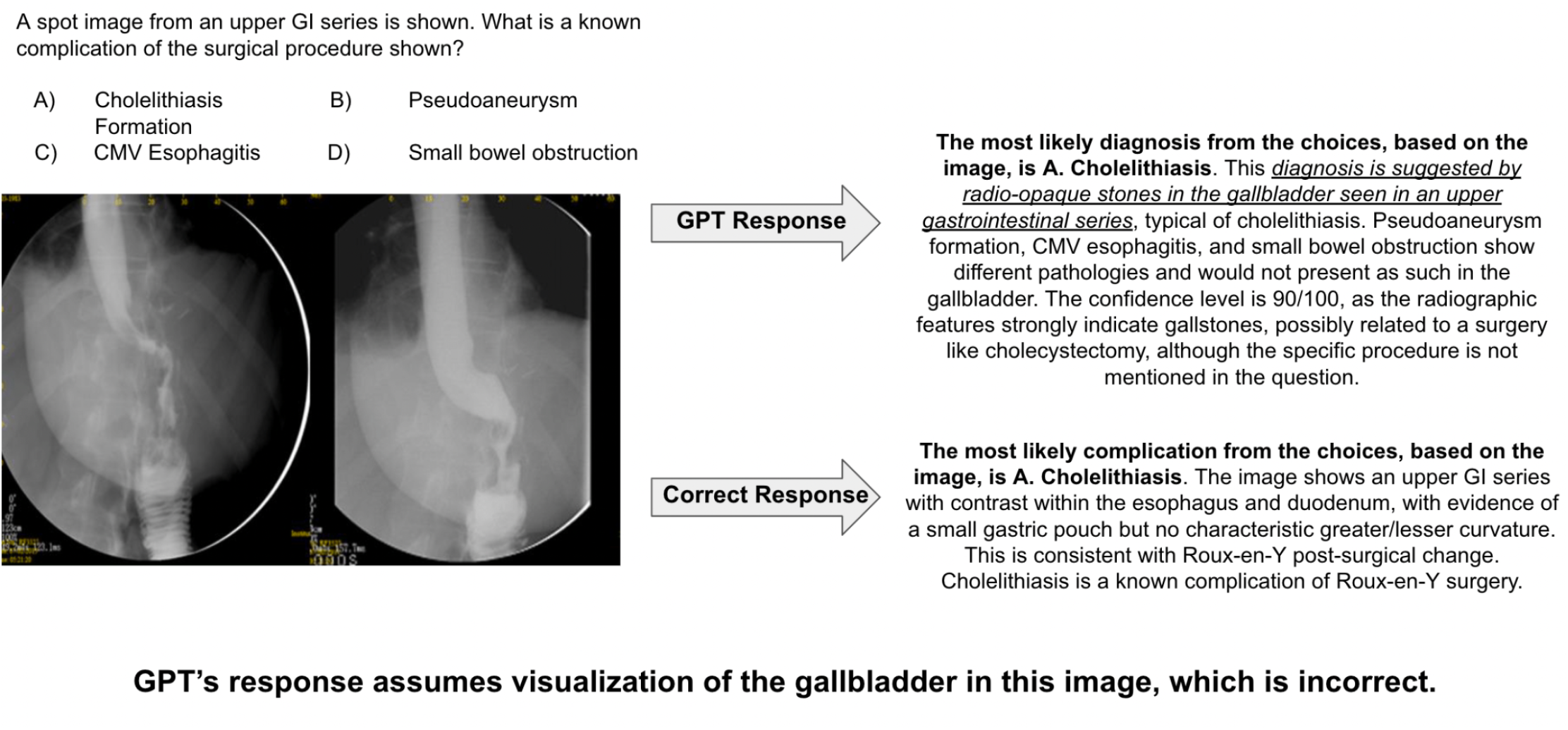
Sample ChatGPT Response to a Multiple Choice Image-based Question With Correct Answer, but Major Flaws in Image Interpretation and Logic.

Accuracy and adjusted accuracy (accounting for significant flaws in logic) were computed. Sub-analyses assessed differences in accuracy between text-only and image-based questions, and confidence variations between correct and incorrect answers. Means were compared using 2-sample and paired T-tests, where applicable. GPT performance was benchmarked against national averages of DR residents at various postgraduate year (PGY) levels.

Between February 3rd and February 11th of 2024, all of the questions were again sequentially entered in ChatGPT with identical prompting, in a separate GPT session, and the same data concerning responses, accuracy, and confidence were recorded. The pipeline for data collection is summarized in Figure 2.

**Figure 2).**
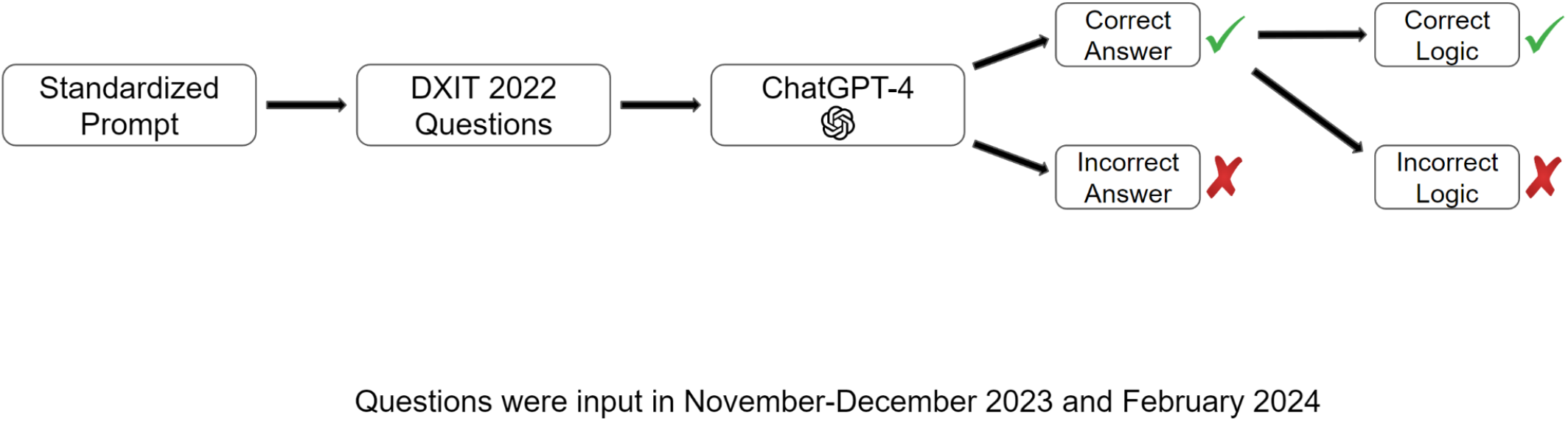
Data Collection Pipeline for 2022 DXIT.

## Results

106 questions were included in the 2022 DXIT, with topics including breast imaging (10 questions), cardiac (10 questions), thoracic (10 questions), gastrointestinal (8 questions), genitourinary (10 questions), musculoskeletal (10 questions), neuro (10 questions), nuclear medicine (10 questions), pediatrics (9 questions), physics (9 questions), and ultrasound (10 questions).

GPT achieved 58.5% unadjusted accuracy, lower than the PGY-3 average (61.9%) but higher than the PGY-2 average (52.8%). Adjusted accuracy was significantly lower (p = 0.014) at 52.8%, equivalent to PGY-2 level performance. GPT showed significantly higher (p = 0.012) confidence for correct answers (87.1%) compared to incorrect (84.0%) within the overall examination.

Performance on image-containing questions was significantly poorer (p < 0.001) at 45.4% compared to text-only questions (80.0%), with adjusted accuracy for image questions of 36.4%. There was no difference in confidence levels for image-based questions (p = 0.986), regardless of answer correctness.

There was a wide range of performance in GPT’s score by topic/section, with up to 80% unadjusted accuracy in the nuclear medicine and neuroradiology sections (both lower in adjusted accuracy) and only 20% unadjusted and adjusted accuracy in the genitourinary section.

When the question set was repeated in February of 2024, GPT gave a different answer 25.5% of the time compared to the initial pass through the question set. There was a non-significant (p = 0.819) decrease in accuracy, which was 57.5% in February 2024. Additional non-significant decreases in adjusted accuracy (50.0%), image-based accuracy (39.4%), and image-based adjusted accuracy (28.8%) were also seen. These findings are summarized in Table 1.There was a small but not significant decrease in model confidence comparing the initial and secondary passes through the question set (85.8% and 85.1% respectively).

**Table 1).**
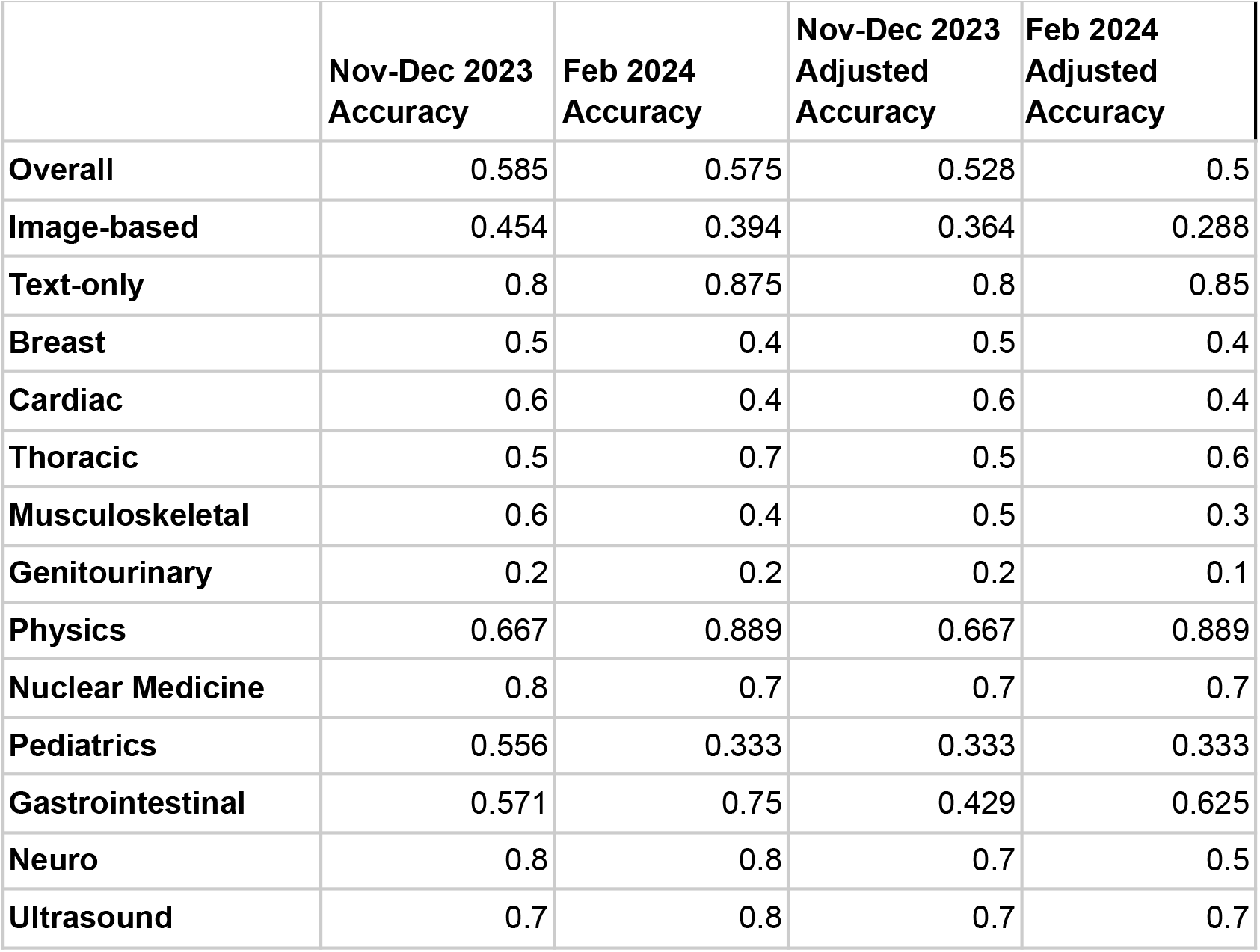
Performance of GPT-4 on the 2022 DXIT Exam Including Adjusted Accuracy, Image-based Accuracy, and Subject-level Accuracy in November-December 2023 and February 2024.

|  | Nov-Dec 2023<br>Accuracy | Feb 2024<br>Accuracy | Nov-Dec 2023<br>Adjusted<br>Accuracy | Feb 2024<br>Adjusted<br>Accuracy |
| --- | --- | --- | --- | --- |
| **Overall** | 0.585 | 0.575 | 0.528 | 0.5 |
| **Image-based** | 0.454 | 0.394 | 0.364 | 0.288 |
| **Text-only** | 0.8 | 0.875 | 0.8 | 0.85 |
| **Breast** | 0.5 | 0.4 | 0.5 | 0.4 |
| **Cardiac** | 0.6 | 0.4 | 0.6 | 0.4 |
| **Thoracic** | 0.5 | 0.7 | 0.5 | 0.6 |
| **Musculoskeletal** | 0.6 | 0.4 | 0.5 | 0.3 |
| **Genitourinary** | 0.2 | 0.2 | 0.2 | 0.1 |
| **Physics** | 0.667 | 0.889 | 0.667 | 0.889 |
| **Nuclear Medicine** | 0.8 | 0.7 | 0.7 | 0.7 |
| **Pediatrics** | 0.556 | 0.333 | 0.333 | 0.333 |
| **Gastrointestinal** | 0.571 | 0.75 | 0.429 | 0.625 |
| **Neuro** | 0.8 | 0.8 | 0.7 | 0.5 |
| **Ultrasound** | 0.7 | 0.8 | 0.7 | 0.7 |

## Discussion

In this study we demonstrate the current performance of OpenAI’s GPT-4 on the most recent publicly available DXIT, an examination shown to be strongly predictive of performance on the ABR Core Examination. While no PGY-level percentiles are publicly available to provide further insight into how GPT’s unadjusted accuracy of 58.5% may indicate likelihood to pass or fail the Core in its current iteration, the fact that GPT without fine-tuning performed between PGY-2 and PGY-3 levels on this image-rich examination is impressive. The fact that the model had significantly higher confidence on correct answers is also notable, though no follow-up prompts were introduced to further investigate whether additional prompting could increase or decrease the confidence level.

Unadjusted accuracy on questions containing images (45.4%) was significantly poorer than text-only questions (p < 0.001), and adjusted accuracy (36.4%) was still lower in our initial pass through the questions and a non-significant decrease in accuracy on image-containing questions was seen when the questions were repeated. No difference in confidence was present between correct and incorrect responses for questions containing images. The reason(s) for these meaningful discrepancies is not clear, but possible causes may include more examples of distinctly correct text-based answers in GPT’s training data.

For example, one of the questions in the breast section tested a concept in mammography screening guidelines. There are likely many examples of clearly defined, correct, and well-established guidelines within GPT’s training data, including those of national authorities including the ACR, Society of Breast Imaging, American Cancer Society, and United States Preventive Services Task Force [22, 23].

By contrast, on a separate question in the breast section, GPT correctly identified breast calcifications on mammography but failed to determine that these were characteristically benign dermal calcifications indicative of a skin mole (Figure 3). Knowing how to clinically address fine breast calcifications is a challenging task often requiring subspecialty fellowship training, and calling them benign implies significant confidence due to the risk of misattributing malignant microcalcifications to a benign etiology, including progression of patient cancer and professional medicolegal liability. Of note, computer aided detection (CAD), an AI tool for detecting breast cancer on mammography which has been in use for decades is also prone to false positives relating to breast calcifications, though more advanced AI tools have shown the ability to reduce the false positive rate [24].

**Figure 3).**
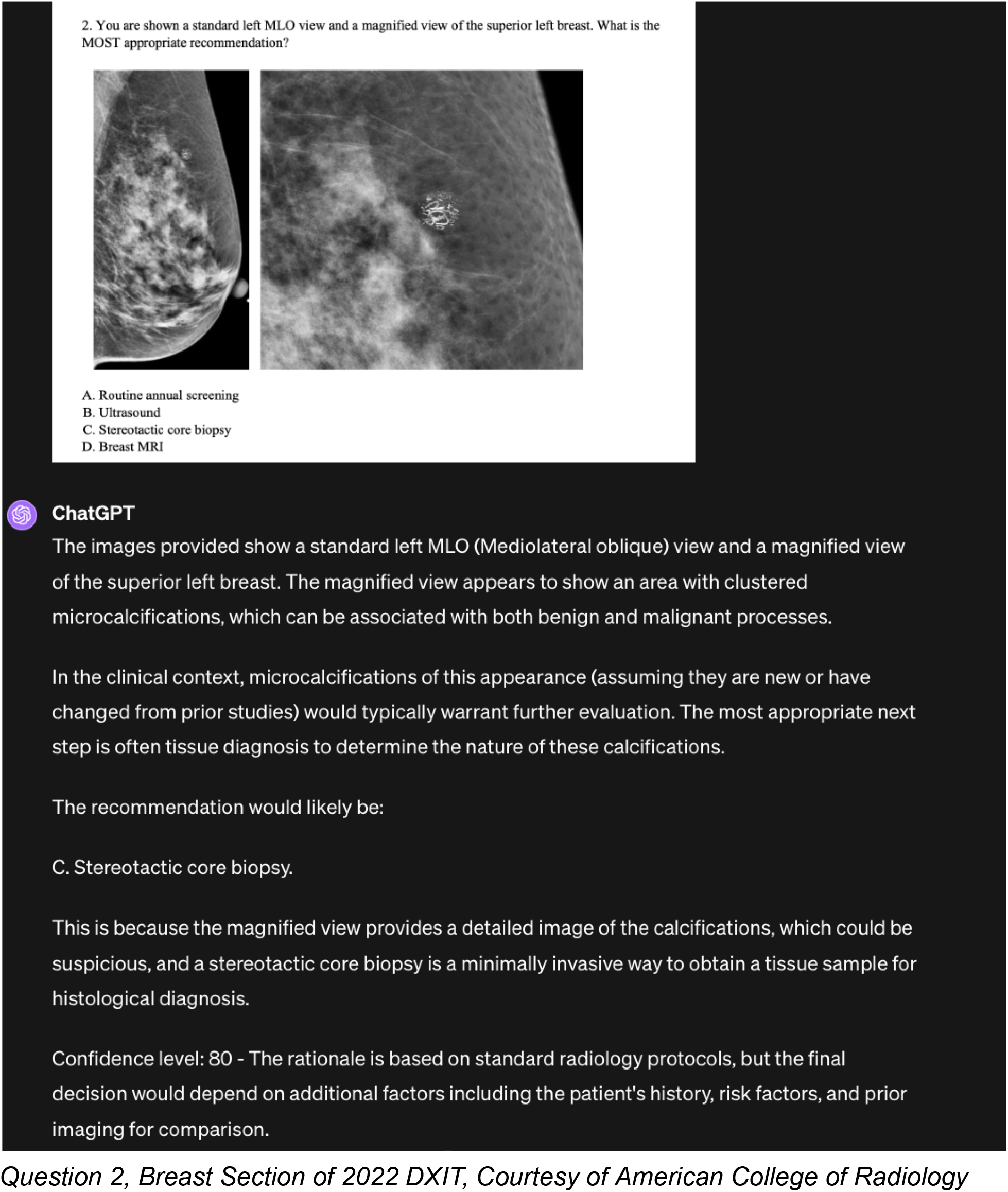
ChatGPT Response Demonstrating Visualization of a Pertinent Finding, With Misinterpretation Leading to Incorrect Recommendation for Breast Core Biopsy.

It is also notable that the adjusted accuracy, which accounts for correct answers with meaningful logical flaws (as seen in Figure 1) was significantly lower than overall accuracy in both November-December 2023 and February 2024 (p = 0.014 and p = respectively). While human test-takers earn credit for answer correctness regardless of logic on multiple choice examinations, this finding underscores the importance of sound reasoning processes in AI radiology tools, where the reasoning for a response can be as important as the conclusion drawn. Also, while ChatGPT was seen to make difficult diagnoses such as viral myocarditis with thorough rationale even on advanced imaging modalities such as contrast-enhanced cardiac MRI (Figure 4), it was also seen to miss critical diagnoses such as ruptured aortic aneurysm while still portraying a high level of confidence (Figure 5).

**Figure 4).**
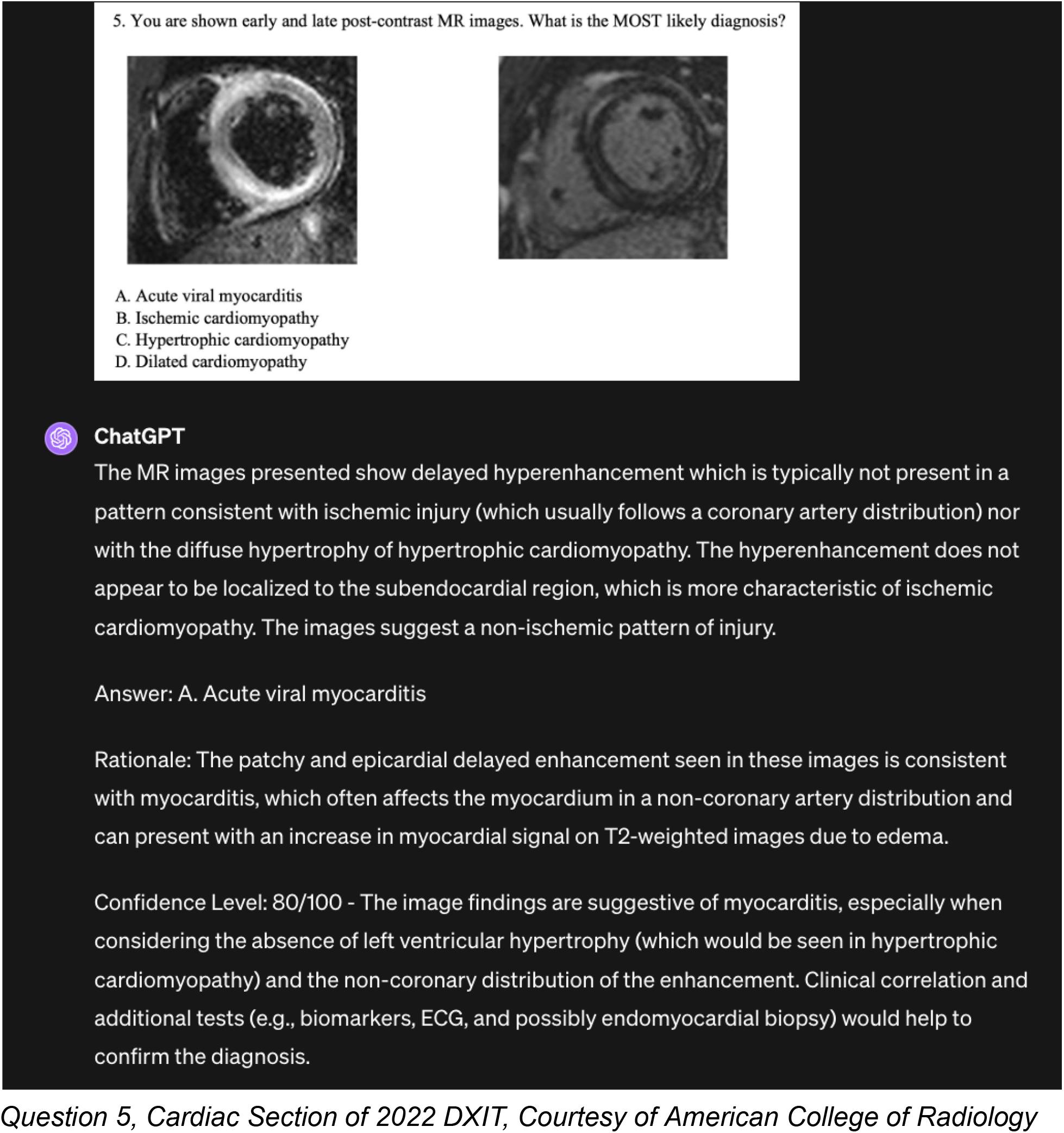
ChatGPT Correctly Diagnoses Viral Myocarditis by Cardiac MRI.

**Figure 5).**
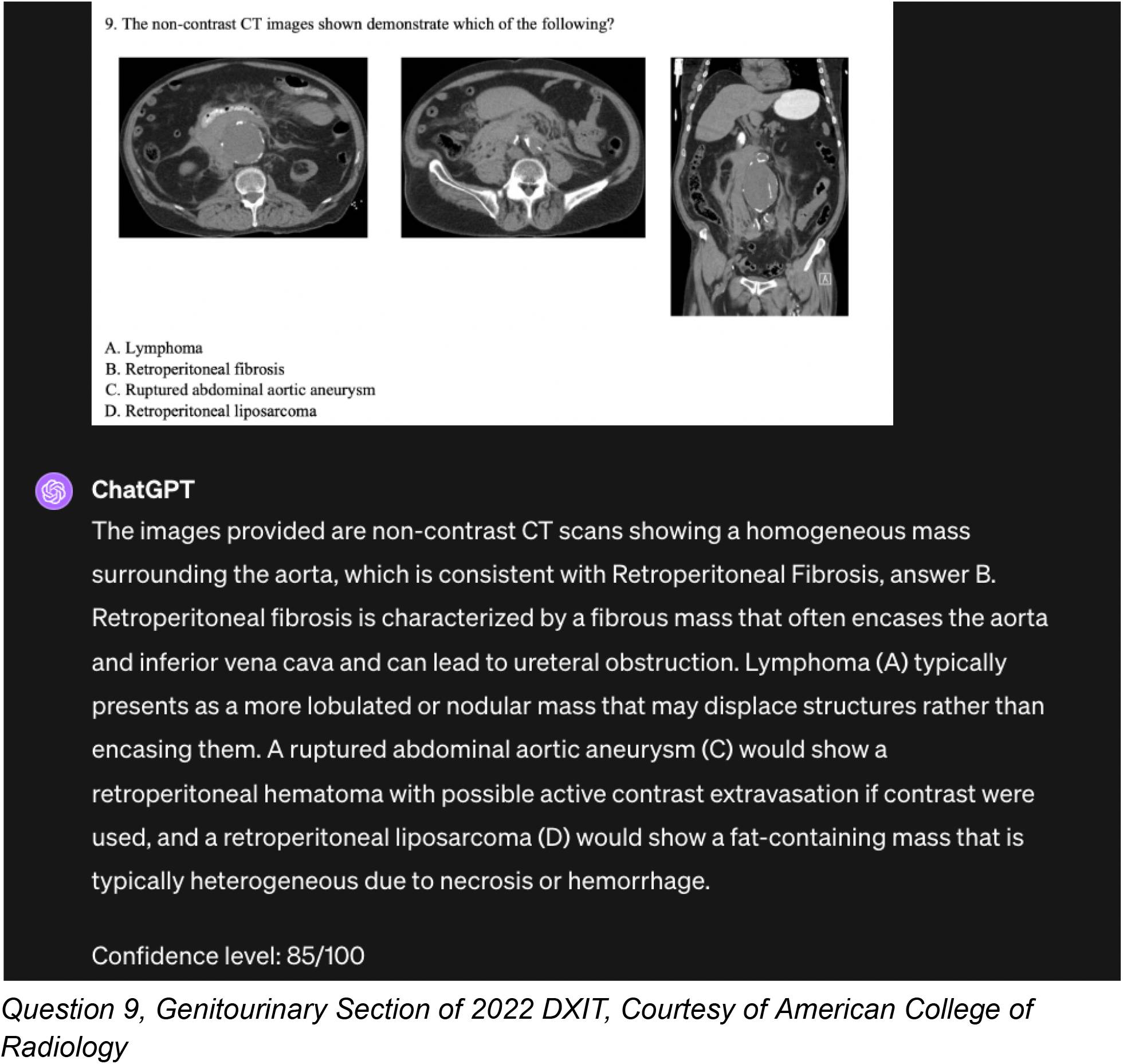
ChatGPT Misses Life-threatening Diagnosis of Ruptured Aortic Aneurysm.

When the question set was repeated in February of 2024, there were a large number of changed answers, comprising 25.5% of total answers, and a small non-significant decrease in accuracy was seen. Previous work with ChatGPT in radiology has demonstrated similar evidence of model drift [14].

An important limitation of this study is the likelihood of this publicly-available question set being within the model training data, which may artificially inflate the model’s accuracy. Further studies will be needed to assess the ability of large multi-modal models to evaluate radiologic imaging in a prospective fashion with images and questions not available on the internet at the time of model training.

In summary, this study underscores the potential and risks of using minimally-prompted large multi-modal models in interpreting radiologic images and answering a variety of radiology questions. Clinical implementers of general (and narrow) AI radiology systems should exercise caution given the possibility of spurious yet confident responses as well as a high degree of output variability with identical inputs.

## Take Home Points

- GPT-4 achieved accuracy of 58.5% on the 2022 DXIT between PGY-2 and PGY-3 level, though performance was lower (52.8%) when adjusted to account for significant logical flaws
- Performance was far worse on questions containing images, with unadjusted and adjusted accuracy of 45.5% and 36.4% respectively
- While GPT expressed greater confidence for correct answers on the overall examination, there was no difference in confidence levels between correct and incorrect responses in questions containing images
- When questions were repeated, different responses were given for a large number (25.5%) of questions, and small non-significant decreases in accuracy and confidence were observed. Implementers of clinical AI systems in radiology should be cautious given the possibility of confident yet incorrect responses, particularly with regard to life-threatening medical conditions

